# Assessing tiger corridor functionality with landscape genetics and modelling across Terai-Arc Landscape, India

**DOI:** 10.1101/2020.10.24.353789

**Authors:** Suvankar Biswas, Supriya Bhatt, Debanjan Sarkar, Gautam Talukdar, Bivash Pandav, Samrat Mondol

**Author notes:** Corresponding authors: Samrat Mondol, Ph.D., Wildlife Institute of India, Dehradun, India. Email-, Bivash Pandav, Ph.D., Wildlife Institute of India, Dehradun, India.

## Abstract

India led the global tiger conservation initiatives since last decade and has doubled its wild tiger population to 2967 (2603-3346). The survival of these growing populations residing inside the continuously shrinking habitats is a major concern, which can only be tackled through focused landscape-scale conservation planning across five major extant Indian tiger landscapes. The Terai-Arc landscape (TAL) is one of the ‘global priority’ tiger conservation landscapes holding 22% of the country’s wild tigers. We used intensive field-sampling, genetic analyses and GIS modelling to investigate tiger population structure, source-sink dynamics and functionality of the existing corridors across TAL. Genetic analyses with 219 tigers revealed three low, but sigficantly differentiated tiger subpopulations. Overall, we identified Seven source and 10 sink areas in TAL through genetic migrant and gene flow analyses. GIS modelling identified total 19 (10 high, three medium and six low conductance) corridors in this landscape, with 10 being critical to maintain landscape connectivity. We suggest urgent management attention towards 2707 sq. km. non-protected habitat, mitigation measures associated with ongoing linear infrastructure developments and transboundary coordination with Nepal to ensure habitat and genetic connectivity and long-term sustainability of tigers in this globally important landscape.

## Introduction

The tiger (*Panthera tigris*) exemplifies one of the major conservation efforts globally. Once distributed across ~30 present-day nations ranging from Armenia to Indonesia and the Russian Far East to India covering a variety of habitats ^1^, their distribution and numbers have drastically reduced to less than 4000 wild individuals at the begining of the twentieth century largely due to habitat loss and human persecution ^2,3^. Despite country-specific conservation efforts, the wild tiger population size was found to be >3200 individuals in 2010 ^4^, leading to a commitment from the heads of 13 tiger-range countries to double their tiger number by 2022 ^5^. As majority of the remaining tigers persisted as small, and often isolated populations ^2,6^ conservation strategies mostly focused on landscape-based approaches to attain recovery goals where habitat improvement, enhanced protection measures, prey augmentation were targeted ^2,3,7,8^. Several priority landscapes were identified as ‘Tiger Conservation Landscapes’ (TCLs) ^2^, where identification of key source populations and consolidation and improvement of surrounding habitats were emphasized ^9–11^. Given that ~80% of global wild tiger population living in most wide variety of habitats were found in India at that time ^12^, the success of the tiger recovery plan and the future survival of the species was mostly dependent on India’s conservation actions. In 2019, India announced the news of doubling its tiger number since 2006 (population estimate of 1411 (1165-1675) in 2006 to 2967 (2603-3346) in 2018) within its existing habitats ^13^. However, these increasing tiger populations now face enormous challenges from increasing human density, rapid urbanisation, expanding agriculture, aggressive infrastructure development and economic growth ^14,15^, and their future persistance will depend on the balance between the deveopmental demands and conservation requirements.

Managing wide-ranging, terrotorial species like tiger at landscape level requires in-depth understanding of key source populations ^4^, identification of potential tiger habitats surrounding these source populations and ensuring habitat connectivity within the landscape ^9,10^. At local level, measures of population dynamics including changes in recruitment and mortality, differential rates of immigration and emigration across sex and age classes, population turnover rates etc. are essential, whereas at landscape scale most emphasis should be given on habitat connectivity to enhance gene flow and reduce the risks from inbreeding ^16–18^. Both of these are of critical importance in the Indian scenario as most of the available protected land (~5% as per the Wildlife Protection Act 1972) is extremely fragmented ^19^ and maintaining connectivity by identifying and managing the critical corridors will play the key role in future tiger conservation ^4,20^. However, generating such detailed information is challenging, time consuming and resource intensive at any scale. In the Indian subcontinent, long-term ecological studies have already identified priority landscapes for tiger conservation (for example, Western Ghats, central India, north-eastern India, Sundarbans and the alluvial Terai flood plains in the Himalayan foothills) that support high potential tiger densities and relatively larger population sizes ^13^. Earlier studies have helped us to understand source-sink population dynamics at landscape-scale in Western Ghats ^8,11,21^ and central Indian landscapes ^17,22–26^ along with Terai habitats of Nepal ^27,28^, but such information is missing from the Indian part of the Terai-Arc landscape (TAL), where population connectivity and their relationship with currently available habitats is poorly understood.

In this paper, we used a combination of intensive field surveys, non-invasive genetic tools and GIS modeling to assess the tiger population connectivity across the TAL. More specifically, we investigated (1) tiger presence and population structure across this landscape covering protected as well as non-protected habitats; (2) population connectivity in terms of tiger source-sink population dynamics in TAL; and (3) identify the critical corridors that helps maintaining the metapopulation dynamics. We addressed these questions by using 219 genetically identified unique tigers across TAL and details from already identified corridors in this landscape ^29^.

## Methods

### Research permissions and ethical considerations

All required permissions for field survey and faecal sampling were provided by the Forest Departments of Uttarakhand (Permit no: 90/5-6), Uttar Pradesh (Permit no: 1127/23-2-12(G) and 1891/23-2-12) and Bihar (Permit no: Wildlife-589). Due to the non-invasive nature (faecal sample based) of the work, no ethical clearance was required in this study.

### Study area

This study was conducted in the Indian part of the TAL. The TAL is the only prime tiger habitat found along the foothills of Himalayas ^20,29^. Covering approximately 28000 km^2^ of forested habitat in the northern India and southern part of Nepal, this region is one of the most important global tiger conservation landscapes ^20,29^. The Indian TAL is a linear 900 km long and 50 km wide landscape with 15000 km^2^ tiger habitat encompassing the north Indian states of Haryana (westernmost), Himachal Pradesh, Uttarakhand, Uttar Pradesh and Bihar (easternmost) (Fig. 1a). Situated at the Himalayan foothills, the habitat supports tropical moist deciduous forests dominated by Sal (*Shorea robusta*), tall Terai swamp grasslands and permanently moist reed swamps ^30^. This landscape is identified as a ‘Global priority’ tiger conservation landscape (TCL) ^2^ and retains about 22% of India’s wild tiger population ^13^. As recent as 2003, tigers were reported across this entire landscape ^29^ but currently found only in the states of Uttarakhand, Uttar Pradesh and Bihar ^13^.

**Figure 1:**
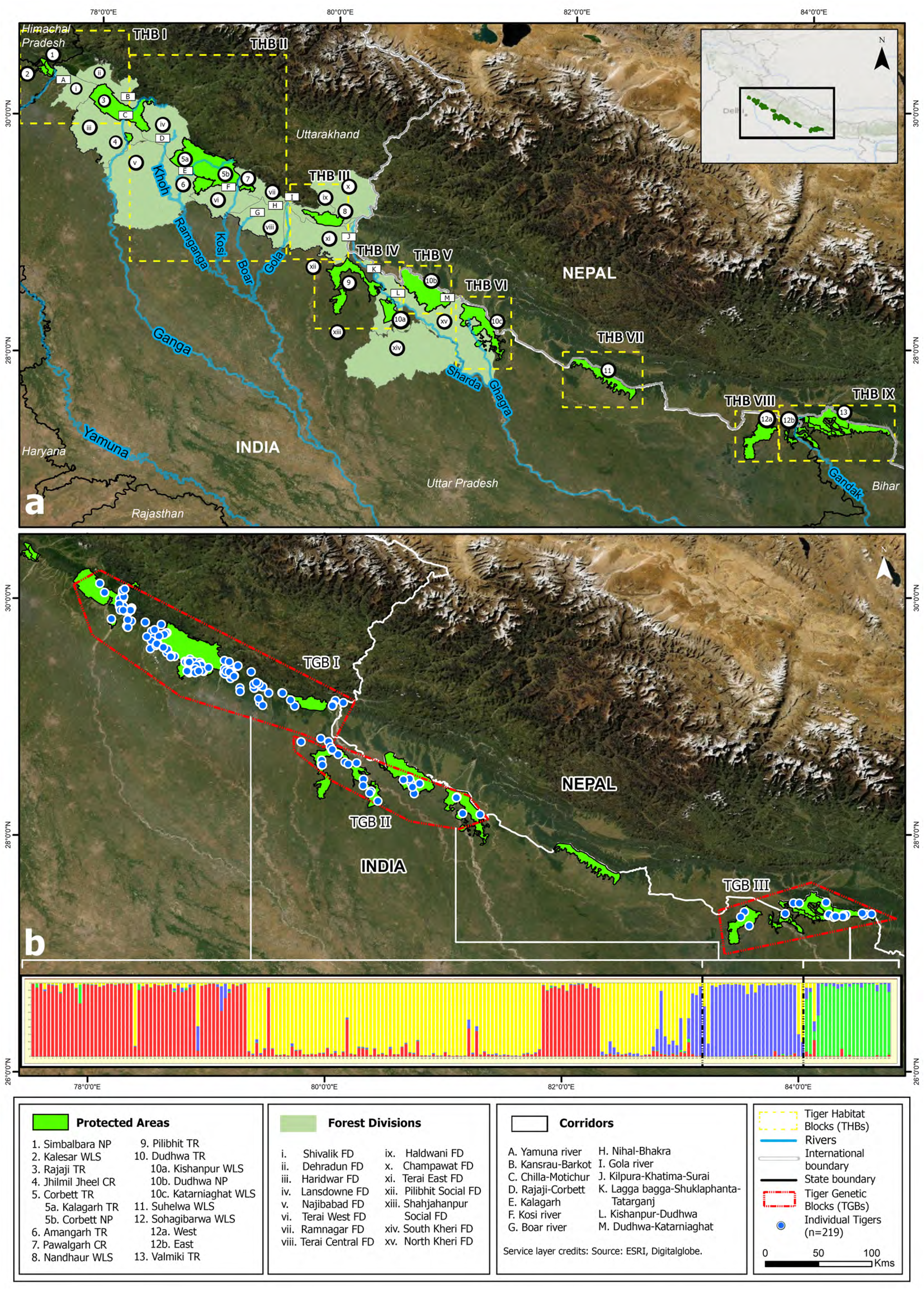
The tiger habiats within the Indian part of Terai-Arc landscape (TAL), encompassing both protected (National parks, Tiger Reserves, Conservation Reserves and Wildlife Sanctuaries) as well as non-protected areas (Forest Divisions and Social Forestry Divisions). The top plot (a) shows the entire landscape with marked ‘Tiger Habitat Blocks’ (THBs) and identified corridors (Johnsingh et al., 2004) ^29^. The bottom plot (b) presents the ‘Tiger Genetic Blocks’ (TGBs) along with the genetic structure results from program TESS ^45^. These TGBs roughly correspond to the western, central and eastern parts of the landscape.

This transboundary tiger habitat in TAL comprises a network of Protected Areas (PAs) and multiple use Forest Divisions (FDs) that maintain habitat connectivity through forest corridors ^20,29^. The first comprehensive landscape-scale study carried out on tiger distribution in Indian TAL by Johnsingh *et al.* (2004) ^29^ highlighted the issue of habitat fragmentation. The study also identified nine tiger habitat blocks (THBs) and 13 structural corridors that potentially facilitate tiger dispersals (Fig. 1a). Currently this entire landscape has 13 PAs (including six Tiger Reserves (TRs), 12 FDs and three Social Forestry Divisions (SFDs)). According to the latest tiger population estimation report ^13^ this landscape hosts 646 (567-726) individual tigers, with a ~33% population increase since the last estimation (n=485 (427-543) ^31^). All relevant details for the THBs are provided in Supplementary Table S1.

### Field sampling

In this study, we aimed to use both direct and indirect approaches (intensive field sampling, genetic analyses and GIS tools) to assess tiger presence, source-sink dynamics and functional connectivity across the entire tiger habitats in TAL. To achieve this goal at such a large landscape scale, it is important to conduct intensive sampling throughout the target study area. We conducted extensive field surveys covering all the THBs (including both PAs as well as FDs) of Uttarakhand, Uttar Pradesh and Bihar between December 2014 to May 2018 (Supplementary Table S1). During field surveys, a team of 5-6 experienced trackers walked this entire region searching for large carnivore faecal samples along all the potential trails and other habitats in this region. Sampling was mostly conducted between November-May i.e. winter and summer seasons every year. The total effort included ~9500 km of foot survey for faecal sampling.

In the field, we categorized large feline carnivore faeces (tiger and leopard) through morphological features such as size, shape, diameter as well as associated field signs (track and scrape marks etc.). We used a dry sampling approach described in Biswas *et al.* (2019) ^32^ for the entire study. Samples were collected with GPS coordinates and kept in dry, dark boxes in the field for a maximum period of one month before they were shipped to the laboratory, where they were stored in −20 °C freezers until further processing. During the sampling period, we collected a total of 1608 relatively fresh large carnivore faecal samples from the study area. The locations of the samples are provided in Supplementary Fig. S1.

### DNA extraction, species, individual and sex identification

We extracted DNA from all the field-collected faeces using protocol described in Biswas *et al.* (2019) ^32^. In brief, we swabbed the top layer of each sample twice with PBS-soaked sterile cotton swabs. Each swab was subsequently lysed overnight at 56 °C with 330 μl ATL buffer and 30 μl of proteinase K (20 mg/ml). Following digestion, swabs were removed and the remaining extraction was followed by the Qiagen DNeasy Tissue Kit (Qiagen Inc., Hilden, Germany) standard protocol. DNA was eluted twice in 100 μl of TE buffer (pH 7.8). For every set of samples (n=22) two extraction negatives were included to monitor contamination.

For species identification, we combined two earlier described tiger-specific molecular markers ^33,34^. The PCR reactions were carried out in 10 μl reaction volumes containing 4 μl of Qiagen master mix (Qiagen Inc., Hilden, Germany), 3 μl of BSA, 1 μl of primer mix and 2 μl of template DNA. The PCR conditions included an initial denaturation (95 °C for 15 min); 50 cycles of denaturation (95 °C for 30 sec); annealing (50 °C for tiger and 57 °C for leopard for 30 sec); extension (72 °C for 30 sec); followed by a final extension (72 °C for 15 min). PCR products were visualized in 2% agarose gels and species-specific band patterns were used for unambiguous species identification. Only tiger samples were subsequently used for individual level analyses.

For individual identification, we used an already standardized panel of 13 microsatellite markers described in Mondol *et al.* (2009) ^35^, Mondol *et al.* (2012) ^36^. These markers were amplified as 10 μl multiplex reactions containing 4 μl of Qiagen multiplex master mix (Qiagen Inc., Hilden, Germany), 3 μl of BSA, 1 μl of primer mix (2 μM concentration) and 3 μl of template DNA. The PCR conditions included an initial denaturation (95 °C for 15 min); 45 cycles of denaturation (94 °C for 30 sec); annealing (Ta for 30 sec); extension (72 °C for 30 sec); followed by a final extension (72 °C for 15 min) in an ABI thermocycler. Post-amplification, 1 μl of PCR product was mixed with 9 μl mix of Hi-Di formamide and 500 LIZ (Applied Biosystems, Foster City, CA, USA) and genotyped in ABI 3500XL DNA fragment analyzer. For genotype validation, we performed a modified multiple tube approach ^37^. Each locus was genotyped and alleles were scored four independent times and a quality index ^35^ was calculated. Only loci with 0.75 or higher quality were retained for further analyses. For allele calling, we expanded the earlier tiger microsatellite allele bins for the north Indian population in program GENEMARKER 2.6.7. (Softgenetics Inc., State College, PA, USA) and scored the alleles manually.

We performed molecular sexing for only individually identified tigers using two sex chromosome markers Amelogenin ^38^ and SRY ^39^. PCR reactions were carried out in 10 μl volumes containing 4 μl of Qiagen master mix (QIAGEN Inc.), 3 μl of BSA, 1 μl of primer mix (3 μM concentration) and 2 μl of template DNA. The PCR conditions included an initial denaturation (95 °C for 15 min); 50 cycles of denaturation (95 °C for 30 sec); annealing (Ta for 30 sec); extension (72 °C for 30 sec), followed by a final extension (72 °C for 15 min) in an ABI thermocycler. PCR products were run in 3% agarose gel and sex identification was done visually through sex-specific banding patterns. For validation, we repeated the entire process twice and only samples provided consensus results were considered further.

### Data analyses

#### Microsatellite summary statistics

We calculated average amplification success as the percent positive PCR for each locus, as described by Broquet and Petit (2004) ^40^. We quantified allelic dropout and false allele (FA) rates manually as the number of allelic dropouts or FAs over the total number of amplifications, respectively ^40^, as well as using MICROCHECKER v 2.2.3. ^41^.

Post data quality assessment, we selected only those samples with good quality data for at least nine loci (out of 13) for further analyses. We identified unique tigers by removing samples with identical genotypes using the ‘Identity analysis’ module of program CERVUS ^42^ and removed the ‘genetic recaptures’ from the dataset. We used the program GIMLET ^43^ to calculate the probability of identity for siblings (PID_(sibs)_) and unbiased (PID_(unbiased)_) for all the individuals. Any allele with >10% frequency across all amplified loci were rechecked and confirmed. We calculated all summary statistics for genetic diversity as well as Hardy Weinberg equilibrium and linkage disequilibrium using program ARLEQUIN 3.1 ^44^.

#### Habitat connectivity in TAL

Assessing tiger connectivity directly (through radio-telemetry or camera trapping) at a landscape level is challenging due to their secretive behaviour, and logistical constraints resulting in infrequent information ^45^. The TAL tiger habitat has 13 identified corridors ^29^ and we aimed to evaluate their current functionality. We took three independent indirect approaches to understand habitat connectivity in TAL. First, we generated individual level tiger genetic information across the entire tiger landscape and used this data to assess any spatial population structure (indicating loss of connectivity). Subsequently, we conducted a number of specific analyses to evaluate source-sink dynamics (using directional gene flow and migrant analyses as proxies of habitat connectivity) involving all the THBs within this landscape to corroborate the population structure results. Finally, we modelled the connectivity between PAs of TAL following Circuit Theory and identified critical corridors to maintain the functionality of this landscape.

For spatial clustering, we performed a Bayesian analysis using program TESS2.3.1 ^46^ with an admixture model with 60000 sweeps and 10000 burn in. We evaluated the most likely number of cluster (K), testing values between 2 to 10 ^47^, where clusters were identified through average deviance information criterion (DIC) value for each K. Further, we calculated genetic differentiation (F_st_ and G_st_) among tiger genetic subpopulations identified through spatial clustering analyses in GenAlEx version 6.5 ^48^.

To assess tiger source-sink dynamics, we performed two different analyses focusing on ‘detection of genetic migrants’ and ‘assessment of rate and direction of gene flow’ within each tiger subpopulation. To detect tiger dispersals, we used two different approaches that use allele frequencies to detect migrant individuals. First, we used prior population information in the USEPOPINFO option implemented in STRUCTURE 2.3.2 ^49^ to detect first-generation migrants in our sampled populations. We detected the number of clusters (K between 1-10) based on 10 independent runs with 500,000 iterations and a burn-in of 50,000 assuming an admixture model. We considered the membership coefficient (*q*) above 0.9 as a realistic cut-off value to assign an individual to a population ^24^. We assigned different migration rates (MIGPRIOR 0.01, 0.05, 0.1) as a sensitivity test. We ran the analysis with two separate dataset: a) individuals grouped as populations according to their sampling locations and b) individuals grouped as genetic clusters from our initial run. These different runs helped us to evaluate the consistency of the results across different genetic groups created.

Further, we used the ‘Migrant detection’ function in program GENECLASS 2.0 ^50^ to identify the first-generation migrant tigers. Here, we used a Bayesian approach as described by Rannala and Mountain (1997) ^51^ along with the resampling method of Paetkau *et al.* (2004) ^52^ for likelihood computation (L_home/L_max). The run parameters included 10000 simulated individuals with a threshold alpha value of 0.01 (Type 1 error) ^52^. This method allows detection of migrants even when the overall differentiation between populations is low. Apart from first generation migrant detection, we did individual assignments in GENECLASS using Bayesian criterion of Rannala and Mountain (1997) ^51^ in combination without resampling with rest of the parameters same as described above. Subsequently, we measured the magnitude and direction of recent gene flow using the Bayesian MCMC based approach implemented in BAYESASS v.3.0.4 ^53^, with run parameters of 1000000 iterations and 100000 burn-in. This approach is generally used to detect recent gene flow (5-7 generations) between populations ^54^.

We developed habitat permeability layer to understand potential tiger connectivity in TAL, using the maximum entropy modelling approach in the program MaxEnt v 3.4.1 ^55^. Habitat permeability layer and focal nodes (regions between which connectivity is to be modelled) are the primary input data for connectivity analyses, indicating the difficulty experienced by an individual in moving across a landscape ^56^. We used tiger presence points and environmental variables in MaxEnt v 3.4.1 ^55^ to develop a habitat permeability layer, where each pixel is a proxy of the likelihood that individuals will move through that cell (i.e. conductance). After removing the spatial cluster between point locations, we used a total 465 presence points along with five environmental variables based on tiger’s ecology and habitat requirements, viz. forest cover, distance to protected areas, distance from water, road, and settlements (Supplementary Table S2). For running the model, we split the presence points as training (70%) and testing (30%) during cross-validate run with 10 replicates and rest of the settings were kept as default. We used the average of all outputs (n=10 runs) as the habitat permeability layer for Circuitscape. Subsequently, we used export to Circuitscape tool (http://www.jennessent.com/arcgis/Circuitscape_Exp.htm) in ArcGIS 10.7 to export the focal nodes and habitat permeability layer to ASCII raster layers of equal extents, cell sizes, and spatial references. We used a total of 21 focal nodes (Supplementary Table S3) and habitat permeability layer in Circuitscape v.4.0 ^56^ to measure the corridor conductance across TAL. We provided weightage to each of the focal nodes based on tiger source-sink populations as identified in this study. We set the node weightages in four categories that ranged from 0 (nodes with no tiger presence), 0.25 (nodes that act as sink populations), 0.5-0.75 (nodes those are moderate-level source populations) and 1 (nodes those are primary source populations) (Supplementary Table S3) during analyses. Subsequently, based on conductance value we classified the output corridors into three categories: low functioning (0-0.30), medium functioning (0.31-0.70) and high functioning (0.71-1.0).

To identify the corridor bottlenecks, we used ‘Pinchpoint mapper’ in Linkage mapper toolbox V2.0 in ArcGIS 10.7. We converted the habitat permeability layer into a resistance surface using SDM toolbox ^57^ in ArcGIS 10.7 (www.esri.com) for running the analysis. Finally, we modelled the least-cost pathways (LCPs) for tiger dispersal using Linkage mapper toolbox V2.0 in ArcGIS 10.7.

## Results

### Tiger data from TAL

Out of 1608 field-collected large carnivore faecal samples, we extracted DNA from 1524 samples (94.78%). Remaining 84 samples (5.22%) showed fungal growth across the top-layer and were not processed. Post tiger-specific assays we identified 743 tiger faeces (48.75%) (Supplementary Fig. S2) for downstream individual identification. Using 13 microsatellite panel ^35,36^ we subsequently identified 219 unique tiger individuals across TAL (Fig. 1b), covering approximately 35% of the estimated tiger population from this landscape (n=646 (567-726) ^13^). Locus-wise mean success rate across all samples ranged between 53-97% (Table 1). We found no large-scale signatures of allele drop out or null alleles. The mean allelic dropout and false allele rates as 0.01 and 0.03, respectively (Table 1), indicating an overall low genotyping error rate across this marker panel. All 13 loci were polymorphic for TAL tigers with mean number of alleles, observed heterozygosity and allelic size range were found to be of 14.76±2.83, 0.35±0.17 and 45.38±10.96, respectively (Table 1). The cumulative PID_(sibs)_ and PID_(unbiased)_ values (2.24*10^−6^ and 5.99*10^−17^, respectively) suggest a statistically strong result for unambiguous tiger individual identification (Table 1). We identified a total of 35 genetic recaptures across TAL. The details of area-wise genetic recaptures are as follows: Rajaji TR- 10, Corbett TR- 11, Pilibhit TR- 2, Lansdowne FD- 5, Hardwar FD- 1, Ramnagar FD- 5 and Terai East FD- 1.

**Table 1:** Genetic diversity and genotyping error details for the individual tigers (n=219) from TAL.

| Locus | Repeat length | N<sub>A</sub> | ASR | H<sub>E</sub> | H<sub>O</sub> | Cumulative PID<sub>(unbiased)</sub> | Cumulative PID<sub>(sibs)</sub> | ADO | FA | Success rate (%) | Reference |
| --- | --- | --- | --- | --- | --- | --- | --- | --- | --- | --- | --- |
| FCA090 | 2 | 19 | 38 | 0.88 | 0.31 | 2.266e-02 | 3.142e-01 | 0.011 | 0.030 | 77 | 35 |
| msFCA506 | 2 | 12 | 38 | 0.85 | 0.26 | 7.991e-04 | 1.045e-01 | 0.003 | 0.050 | 59 | 36 |
| FCA672 | 2 | 15 | 42 | 0.87 | 0.58 | 2.036e-05 | 3.330e-02 | 0.008 | 0.037 | 94 | 35 |
| FCA304 | 2 | 14 | 42 | 0.73 | 0.17 | 2.305e-06 | 1.373e-02 | 0.003 | 0.024 | 92 | 35 |
| FCA628 | 2 | 20 | 54 | 0.83 | 0.41 | 1.007e-07 | 4.721e-03 | 0.012 | 0.026 | 53 | 35 |
| FCA232 | 2 | 16 | 32 | 0.70 | 0.27 | 1.340e-08 | 2.042e-03 | 0.005 | 0.028 | 97 | 35 |
| FCA230 | 2 | 13 | 52 | 0.82 | 0.12 | 7.273e-10 | 7.226e-04 | 0.010 | 0.026 | 94 | 35 |
| FCA279 | 2 | 16 | 36 | 0.87 | 0.51 | 1.761e-11 | 2.298e-04 | 0.000 | 0.031 | 91 | 35 |
| FCA441 | 4 | 14 | 60 | 0.77 | 0.54 | 1.301e-12 | 8.834e-05 | 0.014 | 0.030 | 75 | 35 |
| FCA069 | 2 | 16 | 40 | 0.79 | 0.30 | 7.176e-14 | 3.248e-05 | 0.014 | 0.014 | 79 | 35 |
| msFCA453 | 4 | 12 | 72 | 0.71 | 0.62 | 8.955e-15 | 1.374e-05 | 0.010 | 0.010 | 94 | 36 |
| msF115 | 4 | 09 | 48 | 0.60 | 0.08 | 1.869e-15 | 6.860e-06 | 0.019 | 0.019 | 56 | 36 |
| msHDZ170 | 2 | 16 | 36 | 0.86 | 0.33 | 5.999e-17 | 2.245e-06 | 0.045 | 0.046 | 83 | 36 |
| **Mean** |  | **14.76**<br>**(2.83)** | **45.38**<br>**(10.96)** | **0.79**<br>**(0.08)** | **0.35**<br>**(0.16)** |  |  | **0.012** | **0.028** | **80.3** |  |

N<sub>A</sub>- Number of alleles, ASR- Allelic size range, H<sub>E</sub>- Expected heterozygosity, H<sub>O</sub>- Observed heterozygosity, PID- Probability of identity, ADO- Allelic drop out, FA- False allele

Of all the identified unique individuals (n=219) ~71% were from Uttarakhand (n=156, 35% of the state’s population of 442 (393-491)), whereas ~20% were from Uttar Pradesh (n=44, 25% of the state’s population of 173 (148-198)) and ~9% from Bihar (n=19, 61% of the state’s population of 31 (26-37)) ^13^. We confirmed tiger presence in 12 of the 13 earlier described THBs, representing 10 PAs, nine FDs and two SFDs (Supplementary Fig. S2, Supplementary Table S1). THB VII and part of THB IV did not show any tiger evidence (Supplementary Fig. S2). We achieved an 88% success rate in molecular sexing among the unique tigers (n=193, 89 males and 104 females, respectively). Interestingly, 54 of these males were found inside PAs and 35 individuals were from non-PAs (FDs and SFDs), whereas 59 females were from PAs and 45 from outside PAs, indicating that both sexes were using non-protected habitats across TAL.

### Population structure and genetic differentiation of tigers across TAL

Our Bayesian clustering analyses with 219 individual tiger genetic data revealed four genetic lineages (K=4) across TAL. When examined closely, we found that Uttarakhand state holds two of these four genetic lineages as geographically mixed populations, whereas the remaining two lineages are roughly separated in states of Uttar Pradesh and Bihar, respectively, making three genetic subpopulations across TAL (Fig. 1b). Following the description of ‘Tiger Habitat Blocks (THBs)’ mentioned by Johnsingh *et al.* (2004) ^29^, we report these three genetic subpopulations as ‘Tiger Genetic Blocks (TGBs)’ while reporting the results.

Overall, these TGBs roughly correspond to the tiger population in the states of Uttarakhand (TGB I), Uttar Pradesh (covering mostly TGB II and small parts of TGB I and III) and Bihar (TGB III), respectively (Fig. 1b). The samples collected from THB I, II and III (n=171) formed the first genetic subpopulation (TGB I) (Figs. 1a, 1b). The TGB I is spread between the Rajaji TR (western side) to Nandhaur Wildlife Sanctuary (WLS) (eastern side) in Uttarakhand, covered THB I, II and III and retains the highest tiger population in TAL^13^ (Figs. 1a, 1b). This block represents six PAs, eight FDs, one SFD and nine earlier described corridors (Figs. 1a, 1b, Supplementary Table S1) ^29^. The TGB II includes the samples collected from THB IV, V and VI (n=26) and spread between Pilibhit SFD (western side) to Katarniaghat WLS (eastern side) in Uttar Pradesh (Figs. 1a, 1b). This block retains two PAs, one FD, one SFD and three corridors (Figs. 1a, 1b, Supplementary Table S1) ^29^. Finally, the samples collected from THB VIII and THB IX (n=22) together formed the TGB III and included only two PAs, Sohagibarwa WLS (western side) in eastern Uttar Pradesh and Valmiki TR (eastern side) in Bihar (Figs. 1a, 1b, Supplementary Table S1).

Our analyses revealed that the TGBs are genetically differentiated (Fst and Gst) at low, but significant levels. The Fst value ranged between 0.079-0.107, while the Gst value ranged between 0.067-0.087 (Table 2). Comparative summary statistic analyses among these TGBs showed that TGB I has the highest mean number of alleles (10.15±3.06) when compared to TGB II (7.00±2.11) and TGB III (9.08±2.24), respectively. However, TGB III showed higher observed heterozygosity and the allelic size range values (0.43±0.20 and 38.00±14.29, respectively) than TGB I (0.34±0.20 and 24.46±07.37, respectively) and TGB II (0.29±0.12 and 24.77±09.30, respectively) (Table 3).

**Table 2:** Genetic differentiation (pairwise F_st_ and G_st_) for three TGBs across TAL. The upper diagonal presents pairwise G_st_ values and lower diagonal presents the pairwise F_st_ values

|  | **TGB I** | **TGB II** | **TGB III** |
| --- | --- | --- | --- |
| **TGB I** | 0 | 0.078* | 0.067* |
| **TGB II** | 0.089* | 0 | 0.087* |
| **TGB III** | 0.079* | 0.107* | 0 |

\* p-value= 0.001

**Table 3:** Comparison of different genetic diversity indices among three identified TGBs in TAL.

|  | TGB I (n=173) |  |  |  | TGB II (n=24) |  |  |  | TGB III (n=22) |  |  |  |
| --- | --- | --- | --- | --- | --- | --- | --- | --- | --- | --- | --- | --- |
| Locus | N<sub>A</sub> | ASR | H<sub>E</sub> | H<sub>O</sub> | N<sub>A</sub> | ASR | H<sub>E</sub> | H<sub>O</sub> | N<sub>A</sub> | ASR | H<sub>E</sub> | H<sub>O</sub> |
| FCA090 | 12 | 28 | 0.85 | 0.31 | 08 | 18 | 0.70 | 0.29 | 09 | 28 | 0.78 | 0.37 |
| msFCA506 | 11 | 24 | 0.83 | 0.21 | 08 | 16 | 0.72 | 0.42 | 08 | 34 | 0.84 | 0.38 |
| FCA672 | 13 | 28 | 0.87 | 0.62 | 10 | 24 | 0.88 | 0.27 | 12 | 42 | 0.90 | 0.63 |
| FCA304 | 11 | 28 | 0.73 | 0.14 | 05 | 20 | 0.62 | 0.30 | 05 | 10 | 0.73 | 0.32 |
| FCA628 | 13 | 30 | 0.79 | 0.39 | 09 | 36 | 0.89 | 0.43 | 09 | 54 | 0.79 | 0.59 |
| FCA232 | 11 | 22 | 0.68 | 0.20 | 07 | 20 | 0.53 | 0.43 | 11 | 32 | 0.81 | 0.71 |
| FCA230 | 09 | 26 | 0.80 | 0.10 | 06 | 18 | 0.72 | 0.04 | 10 | 52 | 0.76 | 0.36 |
| FCA279 | 13 | 24 | 0.86 | 0.53 | 09 | 22 | 0.59 | 0.20 | 11 | 32 | 0.77 | 0.73 |
| FCA441 | 11 | 40 | 0.75 | 0.57 | 06 | 36 | 0.74 | 0.31 | 07 | 60 | 0.83 | 0.53 |
| FCA069 | 03 | 26 | 0.71 | 0.33 | 10 | 28 | 0.89 | 0.30 | 09 | 32 | 0.84 | 0.15 |
| msFCA453 | 04 | 08 | 0.65 | 0.70 | 05 | 20 | 0.69 | 0.38 | 10 | 60 | 0.82 | 0.33 |
| msF115 | 10 | 16 | 0.44 | 0.03 | 05 | 48 | 0.74 | 0.35 | 05 | 24 | 0.67 | 0.06 |
| msHDZ170 | 05 | 18 | 0.85 | 0.35 | 03 | 16 | 0.24 | 0.08 | 12 | 34 | 0.90 | 0.36 |
| Mean | 10.15<br>(3.06) | 24.46<br>(7.37) | 0.75<br>(0.11) | 0.34<br>(0.20) | 07.00<br>(2.11) | 24.77<br>(9.30) | 0.69<br>(0.17) | 0.29<br>(0.12) | 9.08<br>(2.24) | 38.00<br>(14.29) | 0.80<br>(0.06) | 0.43<br>(0.20) |

N<sub>A</sub>- Number of alleles, ASR- Allelic size range, H<sub>E</sub>- Expected heterozygosity, H<sub>O</sub>- Observed heterozygosity

### Gene flow and tiger source-sink populations within TGBs

We analysed the genetic data to understand the tiger source-sink population dynamics (which also confirmed connectivity among populations) within each of the three already identified TGBs in TAL. Within the TGB I, ‘genetic migrant detection’ and gene flow analysis identified two major habitat complexes with genetic connectivity: The Rajaji-Lansdowne-Haridwar region and the Corbett-Ramnagar-Terai FDs-Haldwani region. In the Rajaji-Lansdowne-Haridwar habitat complex we identified five first generation migrant tigers (Supplementary Table S4): two migrants each from Lansdowne FD to Rajaji TR and Haridwar FD and one from Rajaji TR to Lansdowne FD (Supplementary Table S4). Further Bayesian analyses show higher rates of gene flow between Lansdowne FD to Rajaji TR and Rajaji TR to Haridwar FD, whereas Najibabad SFD shows signatures of immigration from both Rajaji TR and Lansdowne FD (Fig. 2a). Combining this information, we interpret that both Rajaji TR and Lansdowne FD act as source tiger populations and Haridwar FD and Najibabad SFD are sink populations in this habitat complex, respectively (Fig. 2a).

**Figure 2:**
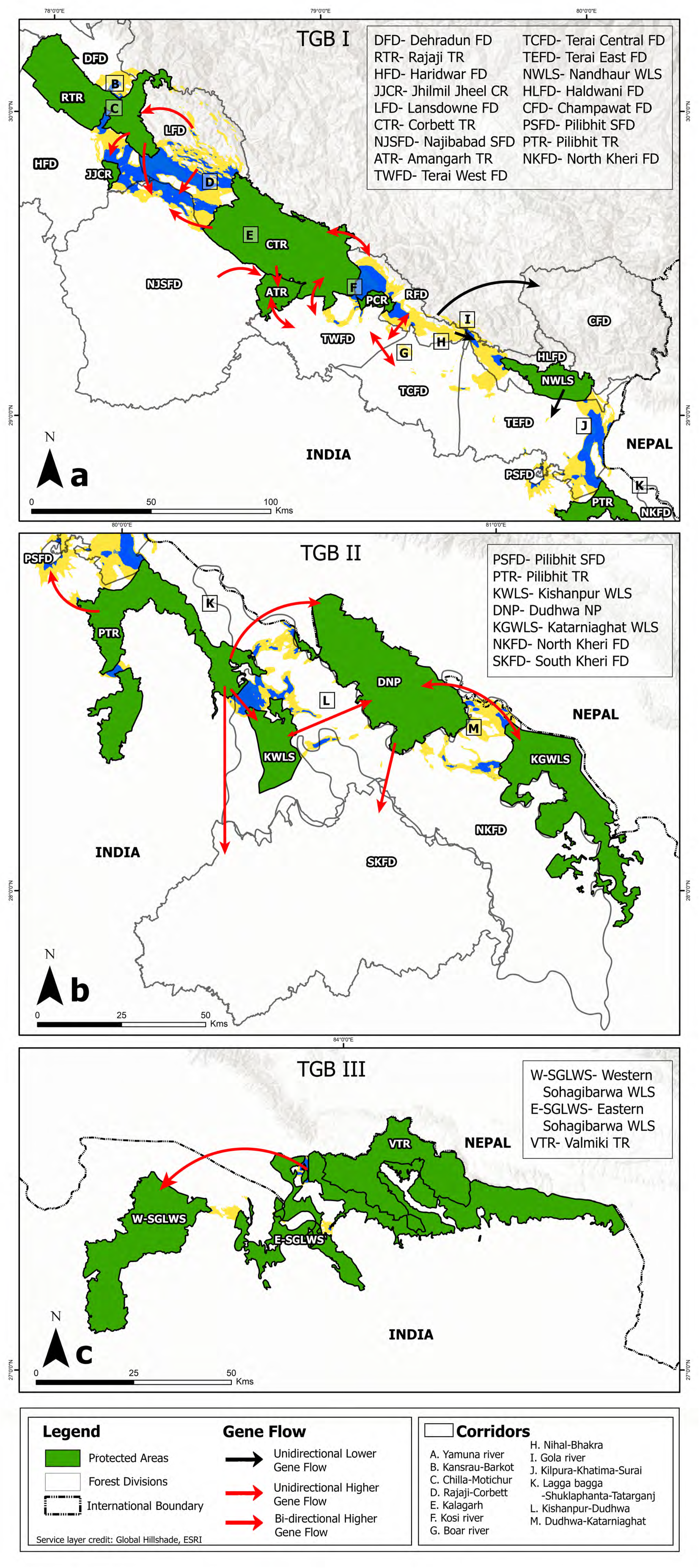
Assessment of tiger source-sink dynamics within each TGB in the Indian part of TAL. The direction and magnitude of gene flow has been presented by different color allows among the protected and non-protected areas. The top plot (a), middlie plot (b) and the bottom plot (c) show the gene flow patterns in TGB I, II and III, respectively.

Similarly, in the Corbett-Ramnagar-Terai FDs-Haldwani habitat complex we identified 12 first-generation migrant tigers (Supplementary Table S4), showing extensive genetic connectivity among the sampled areas within the habitat complex. We found three migrants from Ramnagar FD to Corbett TR and two migrants to Haldwani FD (Supplementary Table S4). We detected one migrant tiger from Terai West FD to Ramnagar FD (Supplementary Table S4). We found two migrants from Terai Central FD to Terai West FD, one migrant to Ramnagar FD and one to Champawat FD (Supplementary Table S4). We also found one migrant individual from Haldwani FD to Terai East FD (Supplementary Table S4). In addition, we detected one migrant individual from Najibabad SFD to Amangarh TR (Supplementary Table S4). Bayesian analyses show higher gene flow from Corbett TR to adjoining Najibabad SFD and Amangarh TR (Fig. 2a). Corbett TR also shows a high rate of bidirectional gene flow with Ramnagar FD and Terai West FD. Similarly, Terai West FD is highly interconnected with Amangarh TR, Terai Central FD and Ramnagar FD (Fig. 2a). The other areas such as Haldwani FD, Ramnagar FD, Terai Central FD and Terai East FD have lower rates of gene flow among them (Fig. 2a). Combined together, this information suggests that Corbett TR is the main source population for Ramnagar FD, Amangarh TR, Terai West FD and Najibabad SFD, whereas Ramnagar FD is the source tiger population for Haldwani, Champawat and Terai East FDs (Fig. 2a). Within the two major tiger habitat complex in TGB I (Rajaji-Lansdowne-Haridwar and Corbett-Ramnagar-Terai FDs-Haldwani) connectivity is maintiained through the Najibabad SFD, which acts as a sink population for both Corbett TR and Lansdowne FD (Fig. 2a, Supplementary Table S4).

Within TGB II, the tiger habitats are comparatively more fragmented than TGB I and we identified 10 first-generation migrant tigers (Supplementary Table S4). We detected two migrants from Pilibhit TR to Dudhwa National Park (NP) (part of Dudhwa TR) and one migrant to Kishanpur WLS (part of Dudhwa TR) (Supplementary Table S4). We also found two migrants from Dudhwa NP to Pilibhit TR, one migrant to Kishanpur WLS (part of Dudhwa TR) and two migrants to South Kheri FD (buffer zone of Dudhwa TR) (Supplementary Table S4). Further, we detected two migrants from Kishanpur WLS to Pilibhit SFD (Supplementary Table S4). We measured higher gene flow from Pilibhit TR to Dudhwa NP, Kishanpur WLS, South Kheri FD and Pilibhit SFD (Fig. 2b). We also estimated higher rate of bidirectional gene flow among Dudhwa NP, Kishanpur WLS and Katarniaghat WLS, along with unidirectional gene flow to adjoining South Kheri FD (Fig. 2b). This information suggests Pilibhit TR and Dudhwa TR are the major source tiger populations, and South Kheri FD and Pilibhit SFD are the sink populations in the TGB II.

Finally, within TGB III we detected two first-generation migrants from west Sohagibarwa WLS to Valmiki TR (Supplementary Table S4). This result was surprising as within this habitat block Valmiki TR has the highest tiger population ^13^. However, the Bayesian analysis results indicate higher gene flow from Valmiki TR and east Sohagibarwa WLS to west Sohagibarwa WLS (Fig. 2c). Combined together, we interpret that Valmiki TR-east Sohagibarwa WLS is the source population for west Sohagibarwa WLS in TGB III.

### Corridor connectivity across TAL

Our habitat permeability analyses indicated certain habitat variables such as distance to forest cover and protected areas are the main governing factors of tiger dispersal (Supplementary Table S5). Out of the 13 earlier-described corridors ^29^, twelve showed conductance for tiger dispersal (Fig. 3a, Supplementary Table S6). Towards the western end of TAL Yamuna river corridor did not show any signatures of habitat conductance (Fig. 3a, Supplementary Table S6). The remaining corridors showed high to low conductances (Fig. 3a, Supplementary Table S6). The details of the least-cost pathways (LCP) are provided in Supplementary Table S6.

**Figure 3:**
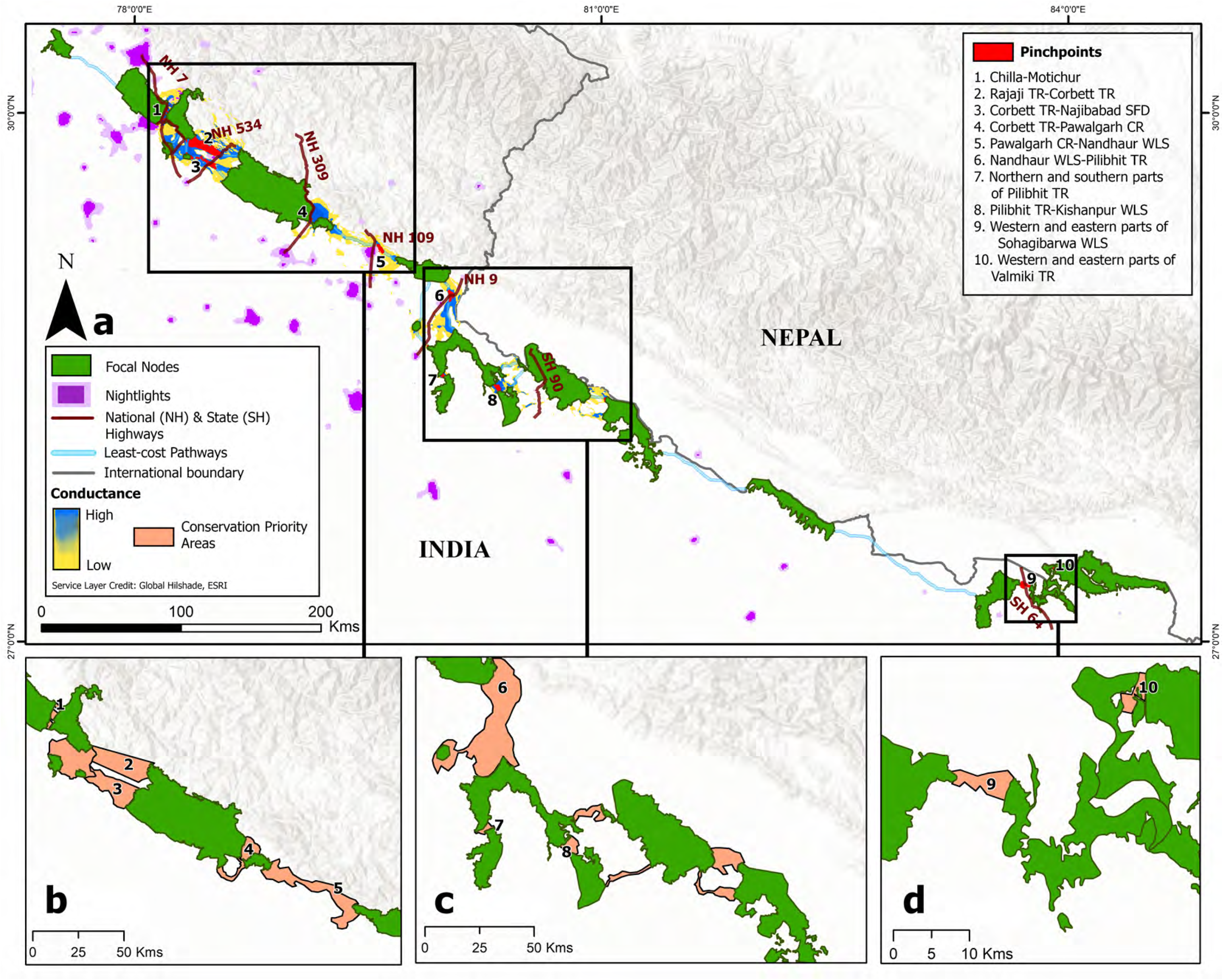
Results of the CIRCUITSCAPE analyses to identify the corridor conductances across TAL. Both ‘Least Cost Pathways (LCPs)’ as well as the critical corridors (pinchpoints) are shown here in (a). The critical corridor areas to maintain contiguous landscape and require urgent management attention are highlighted in (b), (c) and (d). Refer Table 4 for details of these critical areas.

**Table 4:** Details of the critical corridor regions identified in this study.

| SL | Geographic location | Area<br>(sq.km.) | State |
| --- | --- | --- | --- |
| 1 | Connecting western and eastern parts of Rajaji TR<br>(northern block) | 9.73 | Uttarakhand |
| 2 | Connecting western and eastern parts of Rajaji TR<br>(central block) | 14.12 | Uttarakhand |
| 3 | Connecting western and eastern parts of Rajaji TR<br>(southern block) | 9.74 | Uttarakhand |
| 4 | Connecting eastern Rajaji TR, Jhilmil Jheel CR and<br>Najibabad SFD through Haridwar and Lansdowne FDs | 345.07 | Uttarakhand and<br>Uttar Pradesh |
| 5 | Connecting Corbett TR and Najibabad SFD through<br>Lansdowne FD and Najibabad SFD | 267.16 | Uttarakhand and<br>Uttar Pradesh |
| 6 | Connecting Rajaji and Corbett TRs through Lansdowne<br>FD | 328.32 | Uttarakhand |
| 7 | Connecting Corbett TR and Pawalgarh CR through<br>Terai West FD | 53.54 | Uttarakhand |
| 8 | Connecting Corbett TR and Pawalgarh CR through<br>Ramnagar FD | 84.66 | Uttarakhand |
| 9 | Connecting Pawalgarh CR and Nandhaur WLS through<br>Ramnagar, Haldwani and Terai East FDs | 405.37 | Uttarakhand |
| 10 | Connecting Nandhaur WLS and Pilibhit TR through<br>Haldwani and Terai East FDs and Pilibhit SFD | 828.77 | Uttarakhand and<br>Uttar Pradesh |
| 11 | Connecting norther and southern parts of Pilibhit TR | 17.4 | Uttar Pradesh |
| 12 | Connecting Pilibhit TR and Dudhwa NP through North<br>Kheri FD | 56.76 | Uttar Pradesh |
| 13 | Connecting Pilibhit TR and Kishanpur WLS through<br>North Kheri FD | 44.88 | Uttar Pradesh |
| 14 | Connecting Kishanpur WLS and Dudhwa NP through<br>North Kheri FD | 34.83 | Uttar Pradesh |
| 15 | Connecting Dudhwa NP and Katarniaghat WLS<br>through North Kheri FD (northern part) | 115.42 | Uttar Pradesh |
| 16 | Connecting Dudhwa NP and Katarniaghat WLS<br>through North Kheri FD (southern part) | 64.46 | Uttar Pradesh |
| 17 | Connecting western and eastern parts of Sohagibarwa<br>WLS | 18.41 | Uttar Pradesh |
| 18 | Connecting western and eastern parts of Valmiki TR | 8.9 | Bihar |
| **Total** |  | **2707.54** |  |

TR- Tiger Reserve, CR- Conservation Reserve, SFD- Social Forestry Division, FD- Forest Division, WLS- Wildlife Sanctuary, NP- National Park

In addition, we identified seven new corridors in TAL. The first corridor (high conductance) connects Rajaji TR (eastern part) with Najibabad SFD through Lansdowne FD (Fig. 3a, Supplementary Table S6). The second corridor (high conductance) connects Jhilmil Jheel CR with Najibabad SFD through Haridwar FD (Fig. 3a, Supplementary Table S6). The third corridor (high conductance) connects Corbett TR with Najibabad SFD (Fig. 3a, Supplementary Table S6), making these areas very well connected with each other within TGB I. In TGB II, we identified two new corridors (high conductance), where the first one connected the northern and southern part of Pilibhit TR, and the second one connected Pilibhit TR to Kishanpur WLS through North Kheri FD, respectively (Fig. 3a, Supplementary Table S6). Finally, two new corridors were identified in TGB III, the first one (low conductance) connecting western and eastern part of Sohagibarwa WLS and the easternmost corridor (high conductance) connecting two parts of Valmiki TR, respectively (Fig. 3a, Supplementary Table S6). The corridor bottleneck analysis identified 10 critical tiger dispersal corridors in TAL (Fig. 3a, Supplementary Table S6).

## Discussion

We used a multidisciplinary approach to evaluate all tiger corridor functionalities at a landscape level across the TAL. To the best of our knowledge, this is the first study using genetic analyses and corridor modelling to understand tiger source-sink dynamics and connectivity at a landscape level in TAL. We conducted extensive faecal sampling across all tiger habitats and dispersal corridors and confirmed presence of tiger in most of the earlier described THBs in TAL. These areas include already known TRs and FDs ^13^ along with new areas in this landscape (Supplementary Fig. S2, Supplementary Table S1). Our field surveys corroborated with early reports of tiger absence beyond western part of Rajaji TR, Suhelwa WLS, and remaining areas between Dudhwa TR and Sohagibarwa WLS (Fig. 1b, Supplementary Fig. S2) ^13,29^, possibly due to high anthropogenic pressure and habitat loss. Further, we confirmed tiger presence from Jhilmil Jheel Conservation Reserve (CR) (Uttarakhand), Pilibhit SFD and Najibabad SFD (Uttar Pradesh) where they were not reported earlier (Supplementary Fig. S2) ^13^. This expansion in tiger habitat occupancy is possibly due to increasing tiger dispersals to newer areas driven by recent increase in their population in this landscape ^13^. Tigers are territorial and high resource-demanding (in terms of both food and space) animals and such dispersal events are thus normal ^58^. In the Indian subcontinent such dispersal events have been reported from other landscapes also (Northwest India ^59^, Central India ^17,24,25^, Nepal-TAL ^27,28^) and ~35% of the country’s tiger population is reported to live outside the exisiting TRs ^13^. Interestingly, we found that both male and female tigers beyond protected areas including relatively marginal habitats of FDs and SFDs indicating both-sex dispersal events in TAL, as found in other habitats (Central India ^25^, Nepal-TAL ^28^).

Our genetic analyses identified three genetic subpopulations (named as Tiger Genetic Blocks or TGBs) in TAL. This was an unexpected result considering we found tiger presence in majority of the protected and non-protected areas, except some parts of the central and the eastern TAL (Fig. 1b). The easternmost subpopulation TGB III covering Sohagibarwa WLS and Valmiki TR is physically separated from the central and western TAL without any forest connectivity ^29^. Tigers from Valmiki TR have also been reported earlier as genetically unique ^60^. Even in the Nepal part of TAL, three genetic subpopulations have been reported those are connected with TGB II and TGB III ^28^, indicating genetic discontinuity across transboundary sections of TAL in this region. However, the pattern of genetic discontinuity between TGB I and TGB II within the seemingly continuous habitat of central and western TAL was surprising. Similar patterns of tiger population structure has been earlier reported from other tiger landscapes in the subcontinent (for example, Central Indian landscape ^61,25^, Western TAL ^62^, Nepal-TAL ^28^). Even in the same TAL landscape Bhatt *et al.* (2020) ^63^ has reported two leopard genetic subpopulations, possibly driven by landscape features. When compared with this study, it makes ecological sense that TAL leopards showed less population structure than tigers (three TGBs) as they are more habitat generalist and even found in human-dominated areas ^64^. Unlike the other major tiger landscapes in India, both of these sympatric species face one common problem related to this linear landscape where one-dimensional space (resulting in restricted movement opportunities) combined with very high human density makes the entire landscape extremely vulnerable to fragmentation at a relatively small temporal scale ^29^. We strongly feel that the genetic separation between TGB I and II is driven by human-disturbance mediated loss of corridor functionality leading to population differentiation at small, but significant levels (Table 2).

Within each of these TGBs the genetic data helped us to understand the source-sink dynamics and fine-scale population connectivity patterns between the protected and non-protected areas. Our analyses with tiger data from TGB I identified both migrant individuals (n=17) and signatures of high gene flow among majority of the TRs and other surrounding areas (Fig. 2a, Supplementary Table S4), indicating functional corridors within this region. We feel that the complex, undulating forested Shivalik and Bhabhar habitat found in this region has helped in tiger movements as hypothesized by Johnsingh *et al.* (2004) ^29^. Similar pattern was also identified in Western Ghats and central Indian landscapes where tigers were reported to use rough terrain for dispersal and avoided human habitations ^21,65^. In addition, this area also retains the largest number of tigers, mostly within the PAs ^13^ and recent increase in tiger population has probably resulted in migration of the surplus individuals to newer areas. Our source-sink dynamics analyses results also corroborate this pattern, where we found that majority of the sink populations in TGB I are in the flat, Terai region in the southern part (Fig. 2a). The Terai habitat also supports very high human density and known to support poor tiger populations in recent past ^29^. Similarly, Terai habitat dominated TGB II (where majority of the tiger populations are found within PAs) showed good connectivity (Fig. 2b, Supplementary Table S4). Our analyses indicated that mostly the PAs are source population and FD/SFDs are the sink populations (Fig. 2b), showing a contrasting pattern from TGB I where even FDs were source populations (like Lansdowne and Ramnagar FDs) (Fig. 2a). In TGB III, the largest tiger population Valmiki TR was the source whereas the western Sohagibarwa WLS was the only available sink habitat (Fig. 2c).

The Circuitscape analyses provide strong support and possible explanations to the genetic connectivity patterns we observed in this landscape. Overall, we identified a total of 10 high, three medium and six low conductance tiger corridors across TAL (Fig. 3a, Supplementary Table S6), all of them are outside PAs (Fig. 3a). First, the genetic discontinuity between TGB I and II is possibly due to loss of functionality of the Kilpura-Khatima-Surai corridor (marked as corridor J in Fig. 1a) ^29^. This corridor structurally connects the Champawat-Haldwani-Terai East FD complex of TGB I with Pilibhit-Dudhwa TR complex of TGB II but currently dysfunctional (Fig. 1a). Earlier Jonsingh *et al.* (2004) ^29^ reported no tiger use in this corridor, which has been further confirmed by Jhala *et al.* (2020) ^13^. We believe that this loss of connectivity between TGB I and II is possibly resulting from developmental activities (e.g. highway, railway tracks, canal etc.) and habitat loss (from human settlements and urbanization etc.) along the corridor (Fig. 3a). Some recent evidences suggest tiger dispersal events in this corridor ^66^ indicating hope for this corridor but strong management interventions are required to maintain its functionality and ensure one panmictic population between Rajaji to Dudhwa TRs (Fig. 3a). Next, within the TGB I the southern part of Gola river corridor (marked as corridor I in Fig. 1a) connecting Ramnagar-Terai Central FDs complex with Champawat-Haldwani-Terai West FDs complex was identified as a weak corridor (Fig. 3a). While the northern part of this corridor showed medium conductance, the southern part is heavily affected by anthropogenic activities like human settlements, urbanization, industrialization and large-scale boulder mining ^29,67^ leading to low genetic exchange across it (Fig. 2a). Further, we identified that the Chilla-Motichur corridor (marked as corridor C in Fig. 1a) that connects eastern and western parts of Rajaji TR as a weak connecting link (Fig. 3a) ^13,29^. Despite showing high conductance in Circuitscape analysis there is no functional connectivity through this corridor, possibly due to intense anthropogenic activities ^29,68,69^. Further, one of the most important output of this study is the identification of 10 critical bottlenecks distributed across the high, medium and low conductivity corridors (Fig. 3a). Due to the linear geography of the TAL where only one-directional movements are feasible, these corridor bottlenecks demand immediate conservation attentions. Except these identified regions we could not find any other alternative paths (high Himalayan area in north and human-dominated Terai flats in south) and thus maintaining connectivity through these identified corridors are the only way these tiger populations can survive in near future ^67^.

Finally, another important aspect of the results from this study is that despite having a large tiger number and relatively connected populations the TAL tigers have relatively low genetic variation compared to other tiger landscapes within India and Nepal-TAL. Different measures of genetic diversity in TAL showed lower values when compared with central India (N_A_- 12.43 ± 2.99 ^61^, 11.71 ^24^, 9.1 ± 2.2 ^25^; H_O_- 0.65 ± 0.09 ^61^, 0.54 ^24^, 0.70 ± 0.06 ^25^ and ASR- 28.8 ± 10.5 ^25^) and Western Ghats (N_A_- 3-12 ^34^ and H_O_- 0.33-0.68 ^34^) as well as with earlier studies in TAL (N_A_- 6.69 and H_O_- 0.50 ^62^) and Nepal-TAL (N_A_- 3.0 ± 0.33-4.0 ± 0.32 and H_O_- 0.46 ± 0.08-0.58 ± 0.06 ^28^). While such patterns can be possible due to different panel of markers, the same markers have shown higher genetic variation in central India ^24^ and Western Ghats ^34,70^. Interestingly, similar pattern of lower genetic variation (compared to other regions) has also been reported in leopards from TAL, which was attributed to severe population decline in recent years ^63^. TAL has a long history of extensive trophy and bounty hunting of tigers and leopards since Mughal and British era ^71^, possibly leading to a small founder population. Though the current population has increased in recent years ^13^ the small founder population is probably the reason behind lower genetic variations. However, TGB I showed higher genetic variations (N_A(TGBI)_- 10.15 ± 3.06, H_O(TGBI)_- 0.34 ± 0.2 and ASR_(TGBI)_- 24.46 ± 7.37) than TGB II (Table 3), possibly because it hosts the largest tiger population (~500 individuals) in this landscape ^13^. Surprisingly, the small tiger population in TGB III showed relatively higher genetic variation than in TGB II (Table 3). This is possibly due to gene flow from larger tiger populations of Chitwan NP and Parsa Wildlife Reserve of Nepal ^27^, which helped to retain high genetic diversity in the TGB III. The genetic varation of TGB II can possibly be improved by ensuring tiger connectivity through Kilpura-Khatima-Surai corridor from TGB I along with Shuklaphanta NP and Bardia NP of Nepal ^27^.

## Conservation implications

Considering the linear shape of tiger habitats in TAL, fragmentation has always been a major concern for conservation ^29,72^. Maintaining the integrity of this landscape with very high human density and associated linear developments will remain the most important challenge for long-term persistence of tiger in this globally high priority tiger conservation landscape ^29^. Our results on tiger population structure, connectivity and source-sink dynamics thus have critical management implications specially in the backdrop of recent increase in tiger numbers in TAL. One of the encouraging point is the revealing of only three TGBs compared to the eight distinct THBs described by Johnsingh *et al.* (2004) ^29^, indicating considerable amount genetic exchange within these THBs. However, within the TGBs we identified 10 critical corridors (Fig. 3a, Supplementary Table S6) that require urgent attention failing which integrity of TAL will be severely compromised. Further analyses show that a total of 2707 sq. km. habitat (Fig. 3b, 3c, 3d, Table 4) needs appropriate conservation planning to ensure panmictic tiger populations in TAL. Appropriate mitigation measures associated with ongoing linear infrastructure developments, roads in particular, are a must in this highly sensitive conservation landscape ^69^. In addition, TGB II and III requires transboundary cooperation between India and Nepal to ensure maintenance of genetic variation and source-sink dynamics in this population.

We also anticipate another potentially important upcoming tiger conservation challenge in the form of increasing tiger-human interactions. Our results show that significant proportion of tigers in TAL are found outside PAs (Supplementary Fig. S2) (possibly driven by recent increase in tiger population), co-existing with significant amount of human density and associated livestock ^13^. It is likely that the incidences of tiger-human conflict will increase around the sink habitats and corridors in coming years and active management of such conflict situations will be critical for these tigers living in marginalized habitats. Further, adequate attention towards habitat and prey management in the FDs and SFDs where large number of tigers are found (at least in TGB I and II) will play important role in maintaining connectivity of the entire landscape.

India played a leading role in tiger conservation and has achieved a rare global success in population recovery of a large-bodied, apex carnivore ^13^. However, the future of these tiger populations depend on appropriate management of the ever-shrinking habitats and maintaining the existing populations as metapopulations. Our results from TAL showed the functionality of the existing corridors and point out critical areas where immediate conservation attention is needed. We believe that a focused approach to address such concerns will improve the long-term sustainability of the tiger populations in TAL.

## Acknowledgement

We acknowledge the Forest Departments of Uttarakhand, Uttar Pradesh and Bihar for providing necessary permits to carry out the research. Our thanks to the Forest Department officials and frontline staffs for their support and assistance during field surveys. We acknowledge help from Annu, Bura, Abbhi, Ranjhu, Ammi, Inam, Imam, Shrutarshi, Shiv, Tista, Shrushti, Prajak, Harshvardhan, Lakhshminarayan, Ankit, Rakesh, Sultan and Nimisha for their help during field surveys. We appreciate technical help from Tista, Zenab, Aamir and Madhanraj (laboratory work) and other lab members. We thank the Director, Dean and Nodal Officer of Wildlife Forensics and Conservation Genetics Cell of Wildlife Institute of India for their support.

This research was funded by Wildlife Conservation Trust-Panthera Global Cat Alliance Grants and Department of Science and Technology, Government of India (grant no EMR/2014/000982). Samrat Mondol was supported by the Department of Science and Technology INSPIRE Faculty Award (No.IFA12-LSBM-47). The funders had no role in study design, data collection and analysis, decision to publish, or preparation of the manuscript.

## Author contribution

S Mondol and B Pandav conceived the study idea, generated funds and supervised the study. S Biswas and S Bhatt performed the sampling and data generation. Data was analysed by S Biswas (molecular and GIS data), D Sarkar (GIS), G Talukdar (GIS) and S Mondol (molecular data). S Mondol, S Biswas and B Pandav wrote the initial manuscript and all authors approved the final draft.

---

**Supplementary fig. 1:**
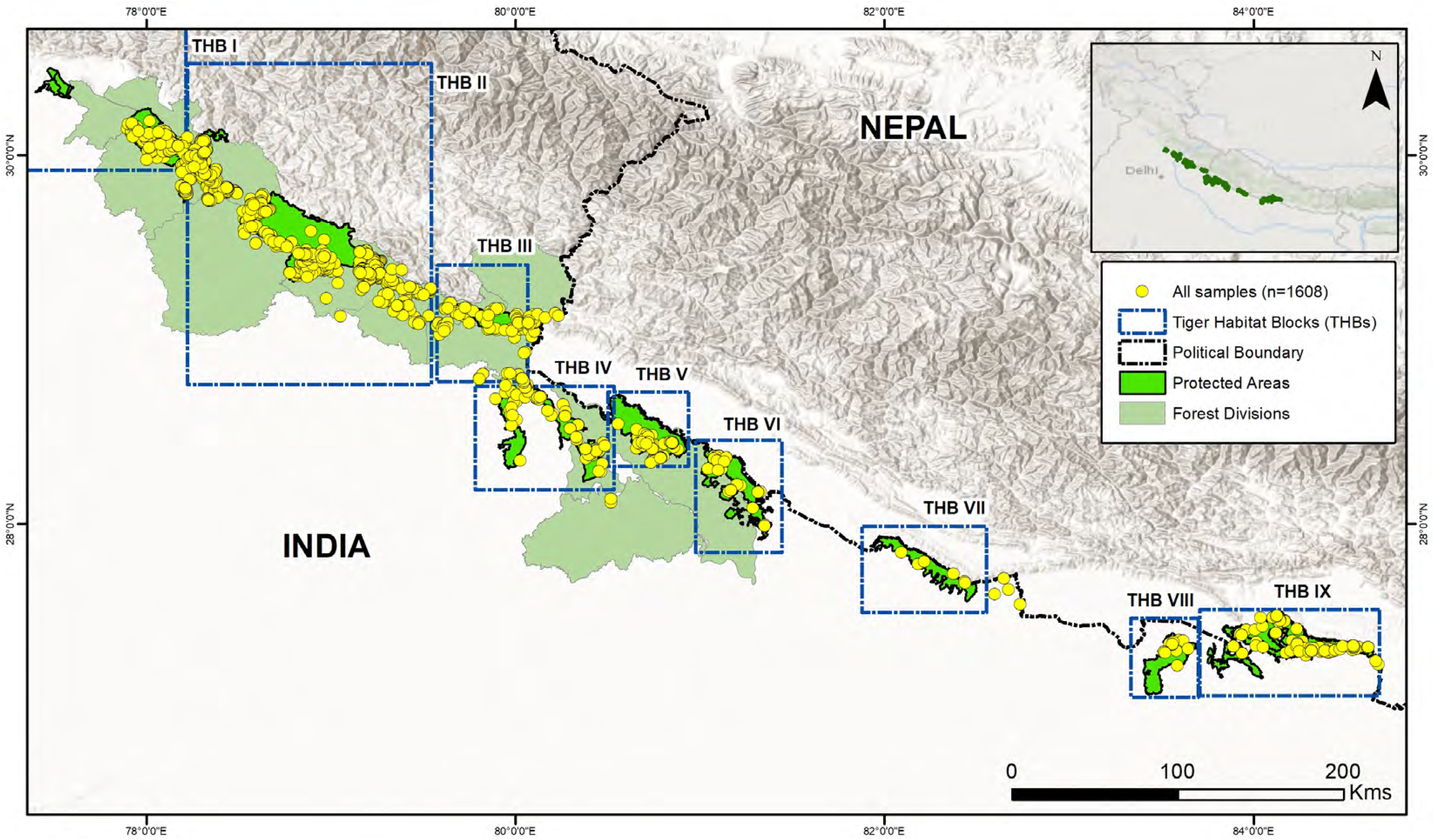
The locations of large carnivore faeces across TAL.

**Supplementary fig. 2:**
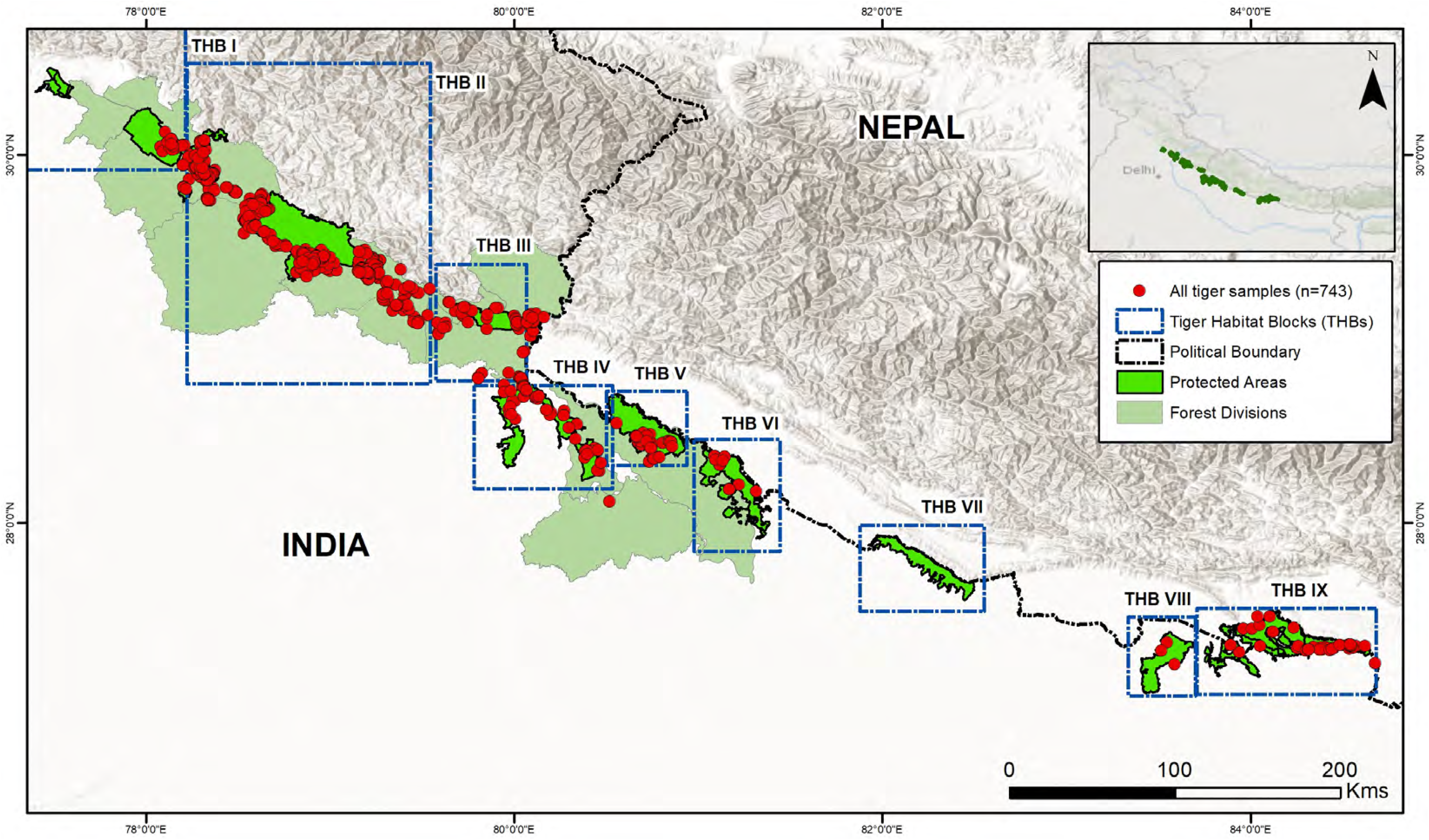
The locations of genetically identified tiger faecal samples across TAL.

**Supplementary Table S1:**
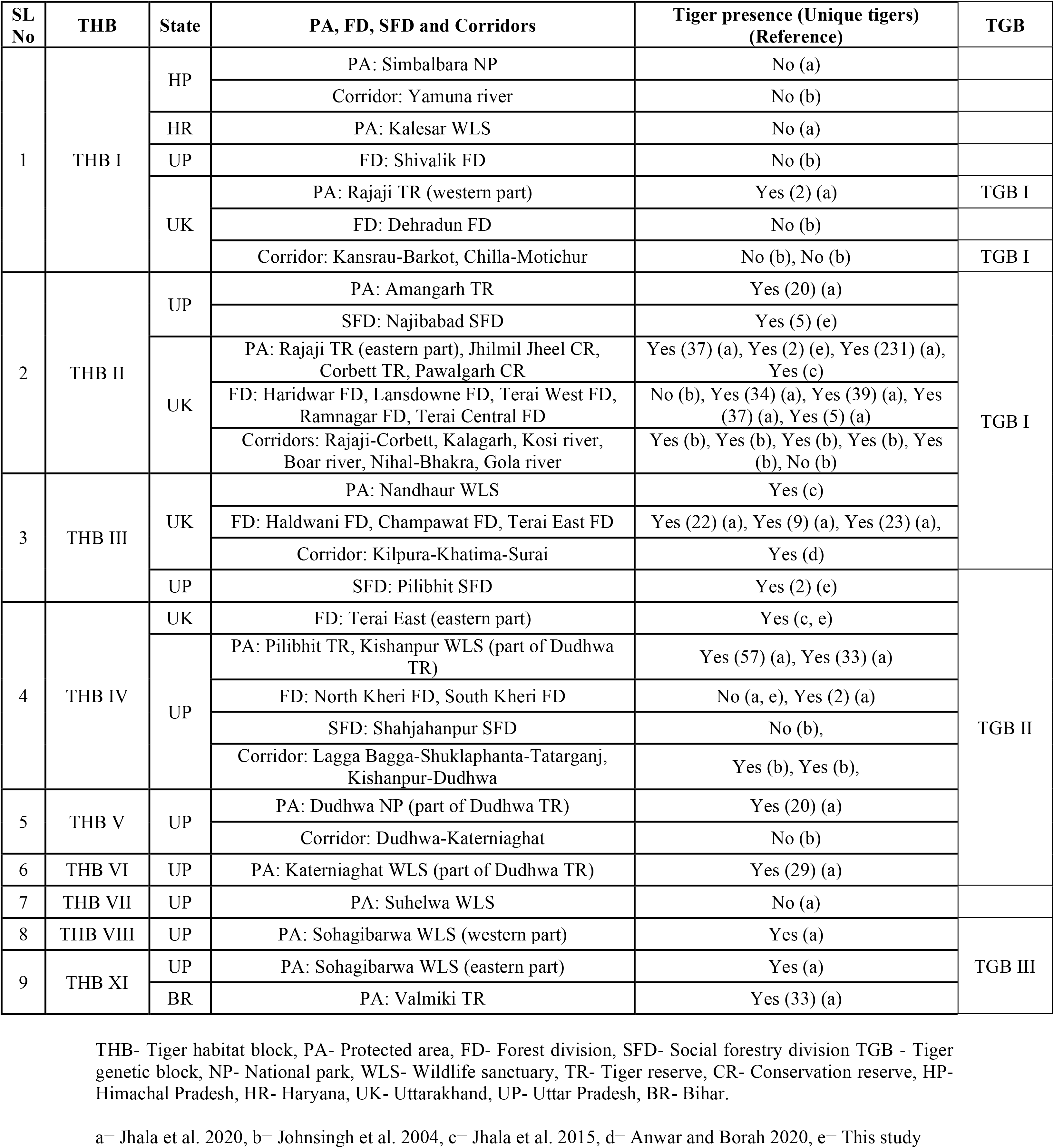
Information of tiger populations, including habitat block details, state, protection status, corridors and identified genetic subpopulations (TGBs) across TAL.

**Supplementary Table S2:** Details of habitat variables used to model tiger habitat permeability in TAL.

| Source file | Covariates used | Source | Unit |
| --- | --- | --- | --- |
| Forest cover | Distance from open/dense forest | Forest survey of India | Meter |
| Protected area (PA) | Distance from PA | World database of protected areas | Meter |
| Road network | Distance from road | Diva-GIS | Meter |
| Settlement | Distance from settlement | Columbia-Village data | Meter |
| River network | Distance from river | Diva-GIS | Meter |

**Supplementary Table S3:** Details of nodes and their weightages used in Circuitscape analysis.

| SL | Name of the node | Protection status of nodes | Weightage (0-1) |
| --- | --- | --- | --- |
| 1 | Kalesar-Simbalbara complex | Protected area | 0 |
| 2 | Rajaji TR- western part | Protected area | 0.50 |
| 3 | Rajaji TR- eastern part | Protected area | 1 |
| 4 | Jhilmil Jheel CR | Protected area | 0.25 |
| 5 | Najibabad SFD | Non-protected area | 0.25 |
| 6 | Corbett TR | Protected area | 1 |
| 7 | Pawalgarh CR | Protected area | 1 |
| 8 | Nandhaur WLS | Protected area | 0.75 |
| 9 | Pilibhit SFD | Non-protected area | 0.50 |
| 10 | Pilibhit TR | Protected area | 0.75 |
| 11 | Pilibhit TR- southern part | Protected area | 0.25 |
| 12 | Kishanpur WLS | Protected area | 0.75 |
| 13 | Dudhwa NP | Protected area | 0.75 |
| 14 | Katarniaghat Wildlife Sanctuary | Protected area | 0.25 |
| 15 | Suhelwa Wildlife Sanctuary | Protected area | 0 |
| 16 | Sohagibarwa WLS- western part | Protected area | 0.25 |
| 17 | Sohagibarwa WLS- eastern part | Protected area | 0.75 |
| 18 | Sohagibarwa WLS- southern part | Protected area | 0 |
| 19 | Valmiki TR- western part | Protected area | 0.75 |
| 20 | Valmiki TR- central part | Protected area | 0.75 |
| 21 | Valmiki TR- eastern part | Protected area | 0.75 |

TR= Tiger Reserve, CR= Conservation Reserve, SFD= Social Forestry Division, WLS= Wildlife Sanctuary, NP= National Park

**Supplementary Table S4:** Detail of the first-generation migrant tigers identified through GENECLASS and STRUCTURE in this study.

| SL | Sampled location | Migrant from | GENECLASS likelihood<br>computation (L home/L<br>max) | *p* Resident | STRUCTURE<br>USEPOPINFO<br>(MIGPRIOR 0.05) |
| --- | --- | --- | --- | --- | --- |
| **TGB I** |  |  |  |  |  |
| 1 | Rajaji TR | Lansdowne FD | 3.645 | 0.003 | NA |
| 2 | Rajaji TR | Lansdowne FD | 3.695 | 0.004 | NA |
| 3 | Lansdowne FD | Rajaji TR | 8.089 | 0.000 | NA |
| 4 | Haridwar FD | Lansdowne FD | NA | NA | 0.936 |
| 5 | Haridwar FD | Lansdowne FD | NA | NA | 0.937 |
| 6 | Terai West FD | Terai Central FD | 7.816 | 0.001 | NA |
| 7 | Terai West FD | Terai Central FD | 5.653 | 0.006 | NA |
| **8** | **Ramnagar FD** | **Terai West FD** | **6.329** | **0.006** | **0.945** |
| 9 | Ramnagar FD | Terai Central FD | 6.307 | 0.004 | NA |
| **10** | **Haldwani FD** | **Ramnagar FD** | **2.347** | **0.006** | **0.960** |
| 11 | Champawat FD | Terai Central FD | 4.993 | 0.000 | NA |
| 12 | Terai East FD | Haldwani FD | 2.374 | 0.000 | NA |
| 13 | Corbett TR | Ramnagar FD | NA | NA | 0.828 |
| 14 | Corbett TR | Ramnagar FD | NA | NA | 0.820 |
| 15 | Corbett TR | Ramnagar FD | NA | NA | 0.922 |
| 16 | Amangarh TR | Najibabad SFD | NA | NA | 0.995 |
| 17 | Haldwani FD | Ramnagar FD | NA | NA | 0.994 |
| **TGB II** |  |  |  |  |  |
| 18 | Pilibhit TR | Dudhwa NP | 4.293 | 0.008 | NA |
| 19 | Pilibhit TR | Dudhwa NP | 4.259 | 0.010 | NA |
| 20 | Kishanpur WLS | Dudhwa NP | 2.647 | 0.000 | NA |
| **21** | **Katarniaghat WLS** | **Pilibhit TR** | **3.773** | **0.000** | **0.981** |
| 22 | Pilibhit SFD | Kishanpur WLS | NA | NA | 0.985 |
| 23 | Pilibhit SFD | Kishanpur WLS | NA | NA | 0.986 |
| 24 | Dudhwa NP | Pilibhit TR | NA | NA | 0.942 |
| 25 | Dudhwa NP | Pilibhit TR | NA | NA | 0.985 |
| 26 | South Kheri FD | Dudhwa NP | NA | NA | 0.984 |
| 27 | South Kheri FD | Dudhwa NP | NA | NA | 0.989 |

| TGB III |  |  |  |  |  |
| --- | --- | --- | --- | --- | --- |
| 28 | Valmiki TR | Sohagibarwa WLS | 5.204 | 0.001 | NA |
| 29 | Valmiki TR | Sohagibarwa WLS | 2.799 | 0.007 | NA |

TGB= Tiger Genetic Block, TR= Tiger Reserve, NP= National Park, WLS= Wildlife Sanctuary, FD= Forest Division, SFD= Social Forestry Division

**Supplementary Table S5:** Contribution of habitat variables in tiger dispersal across TAL.

| **Environmental variables** | **Contribution (%)** |
| --- | --- |
| Distance from forest | 66.8 |
| Distance from protected areas | 23.8 |
| Distance from settlements | 4.6 |
| Distance from river | 3.3 |
| Distance from road | 1.5 |

**Supplementary Table S6:** Details of least-cost pathways, corridor conductance and critical corridors of tiger dispersal identified across the TAL

| SL | Corridors identified between | Length of the Least-cost pathways (km) | Conductance of corridors | Critical corridors |
| --- | --- | --- | --- | --- |
| 1 | Simbalbara-Kaleser complex and Rajaji TR (western part) | 41.5 | NA | NA |
| 2 | Kansrau (western part of Rajaji TR) and Barkot (Dehradun FD) | No | Low | No |
| 3 | Chilla (eastern part) and Motichur (western part) ranges of Rajaji TR | 3.9 | High | Yes |
| 4 | Rajaji TR (eastern part) and Jhilmil Jheel CR | 9.5 | High | No |
| 5 | Rajaji TR (eastern part) and Najibabad SFD | 9.9 | High | No |
| 6 | Jhilmil Jheel CR and Najibabad SFD | 9.5 | High | No |
| 7 | Rajaji TR (eastern part) and Corbett TR | 36.6 | High | Yes |
| 8 | Corbett TR and Najibabad SFD | 31.2 | High | Yes |
| 9 | Corbett TR and Pawalgarh CR | 1.9 | High | Yes |
| 10 | Terai West and Terai Central FDs | NA | Low | No |
| 11 | Terai Central FD and Ramnagar FDs | NA | Low | No |
| 12 | Pawalgarh CR and Nandhaur WLS | 53.8 | Low | Yes |
| 13 | Nandhaur WLS and Pilibhit SFD | 32.2 | NA | NA |
| 14 | Nandhaur WLS and Pilibhit TR | 38.6 | Medium | Yes |
| 15 | Pilibhit SFD and Pilibhit TR | 14 | NA | NA |
| 16 | Northern and southern parts of Pilibhit TR | 1.4 | High | Yes |
| 17 | Pilibhit TR and Kishanpur WLS | 5.6 | High | Yes |
| 18 | Pilibhit TR and Dudhwa NP | 14.8 | Low | No |
| 19 | Kishanpur WLS and Dudhwa NP | 16.2 | Medium | No |
| 20 | Dudhwa NP and Katarniaghat WLS | 11.5 | Medium | No |
| 21 | Katarniaghat WLS and Suhelwa WLS | 71.2 | NA | NA |
| 22 | Suhelwa WLS and west Sohagibarwa WLS | 120.3 | NA | NA |
| 23 | Western and east Sohagibarwa WLS | 7.4 | Low | Yes |
| 24 | Western and eastern parts of Valmiki TR | 0.5 | High | Yes |

TR- Tiger Reserve, CR- Conservation Reserve, SFD- Social Forestry Division, FD- Forest Division, WLS- Wildlife Sanctuary, NP- National Park

